# Dppa2/4 target chromatin bivalency enabling multi-lineage commitment

**DOI:** 10.1101/832873

**Authors:** Mélanie A. Eckersley-Maslin, Aled Parry, Marloes Blotenburg, Christel Krueger, Valar Nila Roamio Franklin, Stephen J. Clark, Clive S. D’Santos, Wolf Reik

**Author notes:** Oncode Institute, Hubrecht Institute–KNAW, University Medical Center Utrecht, Utrecht 3521 AL, The Netherlands. co-corresponding author: MAE-M and WR.

## Abstract

Bivalent chromatin marks developmental promoters in pluripotent cells, yet their targeting and precise impact on lineage commitment remains unclear. We uncover *Developmental Pluripotency Associated 2* (Dppa2) and *4* (Dppa4) as epigenetic priming factors, establishing chromatin bivalency. Single-cell transcriptomics and differentiation assays reveal Dppa2/4 double knockout embryonic stem cells fail to exit pluripotency and differentiate efficiently. Dppa2/4 associate with COMPASS and Polycomb complexes and are required to recruit and maintain their binding at a subset of developmentally important bivalent promoters which are characterised by low expression and poised RNA polymerase. Consequently, upon Dppa2/4 knockout, these dependent promoters gain DNA methylation and are unable to be activated upon differentiation. Our findings uncover a novel targeting principle for bivalency to developmental promoters, poising them for future lineage specific activation.

## Introduction

Epigenetic priming describes the establishment of a competent epigenetic landscape that facilitates efficient transcriptional responses at a future point in time. This temporal uncoupling of molecular events is especially fitting in the context of early development where genes that are not yet expressed need to avoid permanent silencing prevalent in the peri-implantation embryo. Perhaps the best understood example of epigenetic priming is bivalent chromatin. This co-occurrence of active associated H3K4me3 and repressive associated H3K27me3 histone modifications is catalysed by Mll2, part of the COMPASS complex (*1, 2*) and Ezh2, part of the Polycomb Repressive 2 (PRC2) complex, respectively (reviewed in (*3*)). In pluripotent cells, bivalent chromatin is found at important developmental gene promoters, poising them for future activation or silencing (*4, 5*). However, it is largely unclear how these epigenetically primed states are targeted specifically to these promoters (*3*). Furthermore, it is unknown what effect removal of both H3K4me3 and H3K27me3 at bivalent genes has on development as designing a clean experimental system targeting both without altering the rest of the epigenome is challenging. Consequently, our understanding of the precise regulation and functional importance of bivalent chromatin is lacking (*3*).

Recently, we and others revealed a role for the small heterodimerising nuclear proteins *Developmental Pluripotency Associated 2* (Dppa2) and *4* (Dppa4), in regulating zygotic genome activation (ZGA)-associated transcripts (*6–8*). Excitingly, Dppa2/4 also bind non-ZGA gene promoters, including bivalent promoters (*6, 9–11*), however the significance of this is unknown. Intriguingly, single and double zygotic knockout mice survive early embryogenesis only to succumb to lung and skeletal defects shortly after birth, despite these proteins not being expressed in these, or any other adult somatic tissue, uncoupling when they are present from their developmental phenotype (*12, 13*). Here we provide comprehensive molecular and functional evidence supporting Dppa2/4 as novel epigenetic priming factors. Dppa2/4 are required to maintain both H3K4me3 and H3K27me3 at a set of developmentally important bivalent promoters characterised by low H3K4me3 levels and initiating but not elongating RNA polymerase II. As a consequence of losing bivalency, these genes gain DNA methylation and can no longer be effectively activated during differentiation, providing a plausible molecular explanation of the perplexing phenotype of Dppa2/4 knockout mice, and endorsing Dppa2/4 as key developmental epigenetic priming factors.

## Results

### Dppa2/4 promote transcriptional heterogeneity in pluripotent cells, enabling efficient differentiation

Dppa2/4 are frequently used as markers of pluripotent cells, yet their absence has little effect on expression of pluripotent markers at a bulk level (*6, 12, 13*) (Fig. 1SA). Serum/LIF mouse embryonic stem cells (ESCs) are heterogeneous with individual cells varying in their propensity to either self-renew or embark towards differentiation (reviewed in (*14–16*)). During our previous study (*6*), we observed that Dppa2/4 double knockout (DKO) ESC cultures consistently appeared more ‘pluripotent’ with rounder colonies and less spontaneous differentiating cells (data not shown). To investigate this, we performed single-cell transcriptome analysis of wild type (WT) and Dppa2/4 DKO ESCs. Strikingly, Dppa2/4 DKO cells had higher cell-cell correlation indicative of increased homogeneity (Fig. 1A). Dimensionality reduction using UMAP enabled cells to be ordered by pseudotime from more pluripotent towards differentiation (Fig. S1B-C). Remarkably, Dppa2/4 DKO cells preferentially clustered towards the beginning or ‘more pluripotent’ side of the pseudotime trajectory and had very few cells initiating differentiation (Fig. 1B), commonly seen in WT serum/LIF grown cells (*17*). Together this suggested that Dppa2/4 may facilitate exit from pluripotency and cell fate commitment.

**Figure 1:**
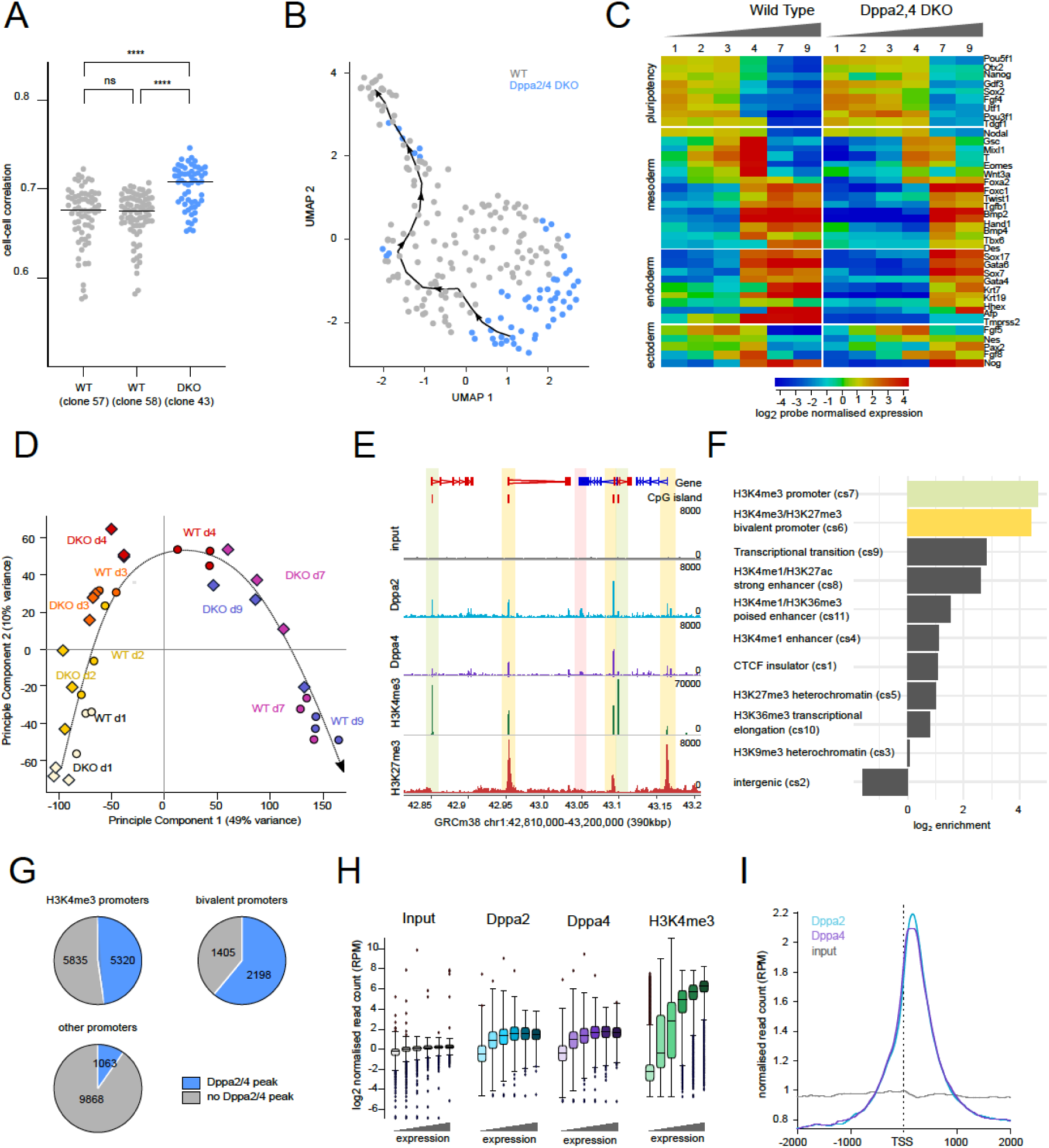
Dppa2/4 are required for differentiation and bind developmental bivalent promoters. (A) Average gene expression correlation values for individual cells to all other cells within wild type (grey) or Dppa2/4 DKO (blue) clones, with higher cell-cell correlation values representing increased gene expression homogeneity. Each dot denotes a single-cell, lines denote mean of distribution, differences are statistically significant (Mann-Whitney Test, **** p-value < 0.0001). (B) Dimensionality reduction (Uniform Manifold Approximation and Projection, UMAP) plot for wild type (WT, grey) and Dppa2/4 DKO (blue) cells. Black line denotes pseudotime from pluripotent (bottom right) towards differentiation (top left). (C) Per gene normalised heatmap showing expression of pluripotency, endoderm, mesoderm and ectoderm genes across 9 days of embryoid body differentiation in WT and Dppa2/4 DKO cells as measured by RNA-sequencing. (D) Principle component analysis of embryoid body time course RNAseq gene expression depicting WT (circles) and Dppa2/4 DKO (diamonds). Biological replicates are denoted by a/b/c. Developmental trajectory is shown by the arrow. (E) Genome browser screenshot showing Dppa2 (blue), Dppa4 (purple) binding and H3K4me3 (green) and H3K27me3 (red) histone modifications across a 394kb region of chromosome 1. Input, Dppa2 and Dppa4 data reanalysed from (*9*), H3K4me3 and H3K27me3 data from this study. H3K4me3 promoters highlighted in green, bivalent promoters highlighted in yellow, inactive promoters highlighted in red. (F) Log2 enrichment of Dppa2/4 peaks amongst 12 chromatin states (cs1-12) defined using ChromHMM models (see Materials and Methods). Chromatin state 12 representing low signal/repetitive elements contained no peaks and is not shown. (G) Proportion of H3K4me3 bound (top left), bivalent (top right) and other (bottom) promoters containing a Dppa2/4 peak (blue). (H) Log_2_ enrichment of Dppa2 (blue) and Dppa4 (purple) binding at gene promoters for different expression bins. H3K4me3 enrichment (green) is shown for comparison. (I) Relative average read density for Dppa2 (blue) and Dppa4 (purple) across a 3kb region centred around the transcription start site (TSS). Input (grey) is shown as a control. Data reanalysed from (*9*).

To test their role in differentiation, we performed embryoid body assays for both WT and DKO ESCs over 9 days from serum/LIF cultures. Importantly, Dppa2/4 DKO cells were delayed in exiting pluripotency and upregulating markers of all three germ layers (Fig. 1C, Fig. S1D-G, Table S1). Supporting this, principal component analysis revealed that the transcriptome of DKO cells at day 7 and 9 of differentiation more closely resembled WT cells at day 4 of differentiation, confirming an overall defect in exiting pluripotency and multi-lineage commitment in these cells (Fig. 1D).

### Dppa2/4 bind H3K4me3 and H3K4me3/H3K27me3 bivalent gene promoters

Bivalent chromatin is a signature of developmental gene promoters in pluripotent cells, however, it is unclear how it is targeted. We, and others, have recently revealed binding of Dppa2/4 at gene promoters in mouse ESCs (*6, 7, 9–11*) (Fig. 1E). Assigning Dppa2/4 peaks to one of 12 chromatin states previously defined by ChromHMM (see Materials and Methods) revealed a dramatic enrichment at both H3K4me3 and H3K4me3/H3K27me3 bivalent promoters (Fig. 1F). Indeed, nearly half of H3K4me3 promoters and over 60% of bivalent promoters defined previously by sequential ChIP-seq experiments (*18*) contained a Dppa2/4 peak, in contrast to just 9.7% of all other protein-coding promoters (Fig. 1G). Stringent peak calling revealed that 22% of Dppa2/4 peaks occur within 1kb of transcription start sites (TSS) (Fig. S1H) of both highly and lowly expressed genes (Fig. 1H), with a clear enrichment of both Dppa2 and Dppa4 at the +1 nucleosome position (Fig. 1I).

### Dppa2/4 interact with members of Polycomb and COMPASS complexes

Dppa2/4 proteins contain both a SAP DNA binding and C-terminal histone binding domain but have no known enzymatic function (*19, 20*). We reasoned that they may instead function by recruiting and/or stabilising enzymatic complexes to their target loci, and therefore performed qPLEX-RIME to quantitatively assess protein interactions with Dppa2/4 in a chromatin context (*21*). We generated independent stable cell lines expressing Dppa2-GFP or Dppa4-GFP, which were expressed at similar levels to the endogenous proteins (Fig. 2A). Importantly, the cells didn’t change expression of pluripotency markers (Fig. S2A) and had minimal changes in gene expression compared to control cells expressing GFP alone (Fig. S2B-C, Table S2), with the largest transcriptional changes occurred at zygotic genome activation (ZGA) associated transcripts (Fig. S2B-D), consistent with additional roles of Dppa2/4 in regulating ZGA-transcription (*6–8*).

**Figure 2:**
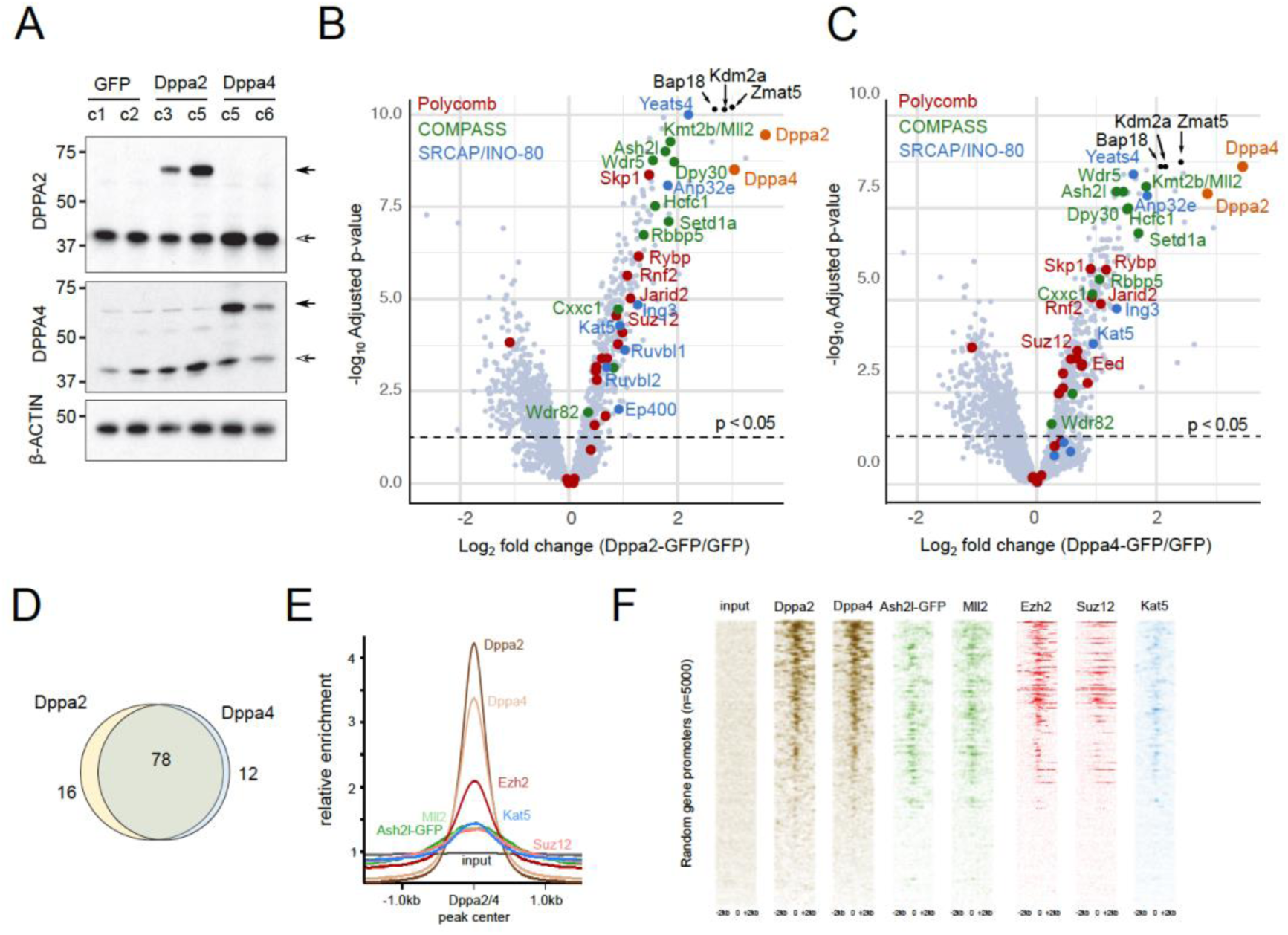
Dppa2/4 associate with members of Polycomb, COMPASS and SRCAP complexes. (A) Western blot showing levels of overexpression of Dppa2-GFP (top) and Dppa4-GFP (middle) in different stable clonal cell lines. β-actin (bottom) is shown as loading control. Open arrows denote endogenous protein, filled arrows overexpressed GFP-tagged protein (B, C) qPLEX-RIME results showing log_2_fold change between Dppa2-GFP (B) or Dppa4-GFP (C) and GFP control versus -log_10_(adjusted p-value). Dotted line represents cut off of p<0.05. Dppa2 and Dppa4 are shown in orange. Top 3 interactors are shown in black. Members of Polycomb (red), COMPASS (green) and SRCAP/INO-80 (blue) complexes are highlighted. (D) overlap between Dppa2 and Dppa4 qPLEX RIME results. (E) Probe trend plot showing relative enrichment across Dppa2/4 peaks. Input, Dppa2 and Dppa4 ChIP data reanalysed from (*9*), Ash2l-GFP data from (*1*), Mll2 data from (*2*), Ezh2 and Suz12 data from (*32*) and Kat5/Tip60 data from (*33*). (F) Aligned probe plots of a random selection of 5000 genes showing enrichment of different chromatin proteins ordered by Dppa4 levels. Data sourced as in (E).

We identified 94 and 90 proteins enriched with Dppa2 and Dppa4 bound chromatin respectively when compared against GFP control (adjusted p-value <0.05, log_2_fold change >1) (Fig. 2B-C, Table S3), of which 78 were in common to both (Fig. 2D). Importantly, mRNA levels of the proteomics hits were not significantly different between WT and DKO cell lines (Fig. S2D). Dppa2 interacted with Dppa4 and *vice versa*, consistent with the proteins functioning as a heterodimer (*13*). Excitingly, Dppa2/4 interacting proteins were enriched for members of both the COMPASS and Polycomb complexes responsible for H3K4me3 and H3K27me3 deposition respectively (Fig. 2B-C). We also detected members in common to the SRCAP and INO80 remodelling complexes, the former of which incorporates the histone variant H2A.Z into nucleosomes. Also of interest were interactions with proteins implicated in zygotic genome activation including Zscan4 and SUMO family proteins (Table S2), consistent with ours and others recent findings that Dppa2/4 also regulate ZGA transcriptional networks (*6–8*).

The association with COMPASS and Polycomb complex members was of particular interest given the localisation of Dppa2/4 at H3K4me3 and bivalent gene promoters. As a validation of our chromatin-based proteomic analysis, we analysed published ChIP-seq datasets from mouse ESCs to verify these complexes bind the same loci as Dppa2/4 (Fig. 2E-F). Importantly, we saw enrichment of COMPASS members Ash2l and Mll2/Kmt2b, Polycomb members Ezh2 and Suz12, and SRCAP member Kat5/Tip60 at Dppa2/4 peaks (Fig. 2E). Moreover, these complex members were also enriched at the same gene promoters as Dppa2/4 (Fig. 2F). Furthermore, we were able to confirm a direct interaction between Dppa4 and Ruvbl1 of the SRCAP/INO80 chromatin remodelling complex (Fig. S2E), consistent with mass spec studies revealing interactions between Dppa2 and Ruvb1, Ruvb2 and Dmap1 (*9*), and Dppa2/4 with H2A.Z (*22*). While we did not reveal a direct strong interaction between Dppa4 and Ash2l or Suz12 in the absence of chromatin (Fig. S2E), others have recently shown a direct interaction between Dppa2 and Suz12 (*9*), suggesting that interactions between Dppa2/4 and the Polycomb and Trithorax machinery are transient and/or substoichiometric and are facilitated by a chromatin template. Therefore, Dppa2/4 bind bivalent gene promoters and interact with the SRCAP/INO-80 chromatin remodelling complex together with the COMPASS and Polycomb complexes in pluripotent cells.

### Loss of both H3K4me3 and H3K27me3 at a subset of bivalent genes in the absence of Dppa2/4

Given their localisation at H3K4me3 marked and bivalent gene promoters and chromatin-based association with COMPASS and Polycomb complex members, we next determined the effect of removing Dppa2/4 on the epigenetic landscape. We first performed ChIP-seq for H3K4me3 and H3K27me3 in WT and Dppa2/4 DKO ESCs. Globally, levels of H3K4me3 and H3K27me3 were largely unchanged (Fig. S3A). Interestingly, 3,179 of 20,897 (15.2%) H3K4me3 and 2,162 of 31,041 (7.0%) H3K27me3 peaks were reduced in Dppa2/4 DKO cells which were predominantly bound by Dppa2/4 and mostly overlapped with gene promoters (Fig. 3A-B). In total, 1,447 and 817 promoters in Dppa2/4 DKO cells had a significant lower enrichment of H3K4me3 and H3K27me3, respectively (Fig. 3C, Table S4). Of these, 611 overlapped, suggesting that bivalent domains may be affected. We therefore focused our analyses on a set of high confidence bivalent promoters defined by ChIP-reChIP experiments in ESCs (*18*). From this, we identified a subset of 309 (9.6%) out of 3,208 bivalent promoters that had lost both H3K4me3 and H3K27me3, and another 327 (10.2%) that had lost just H3K4me3 (Fig. 3C).

**Figure 3:**
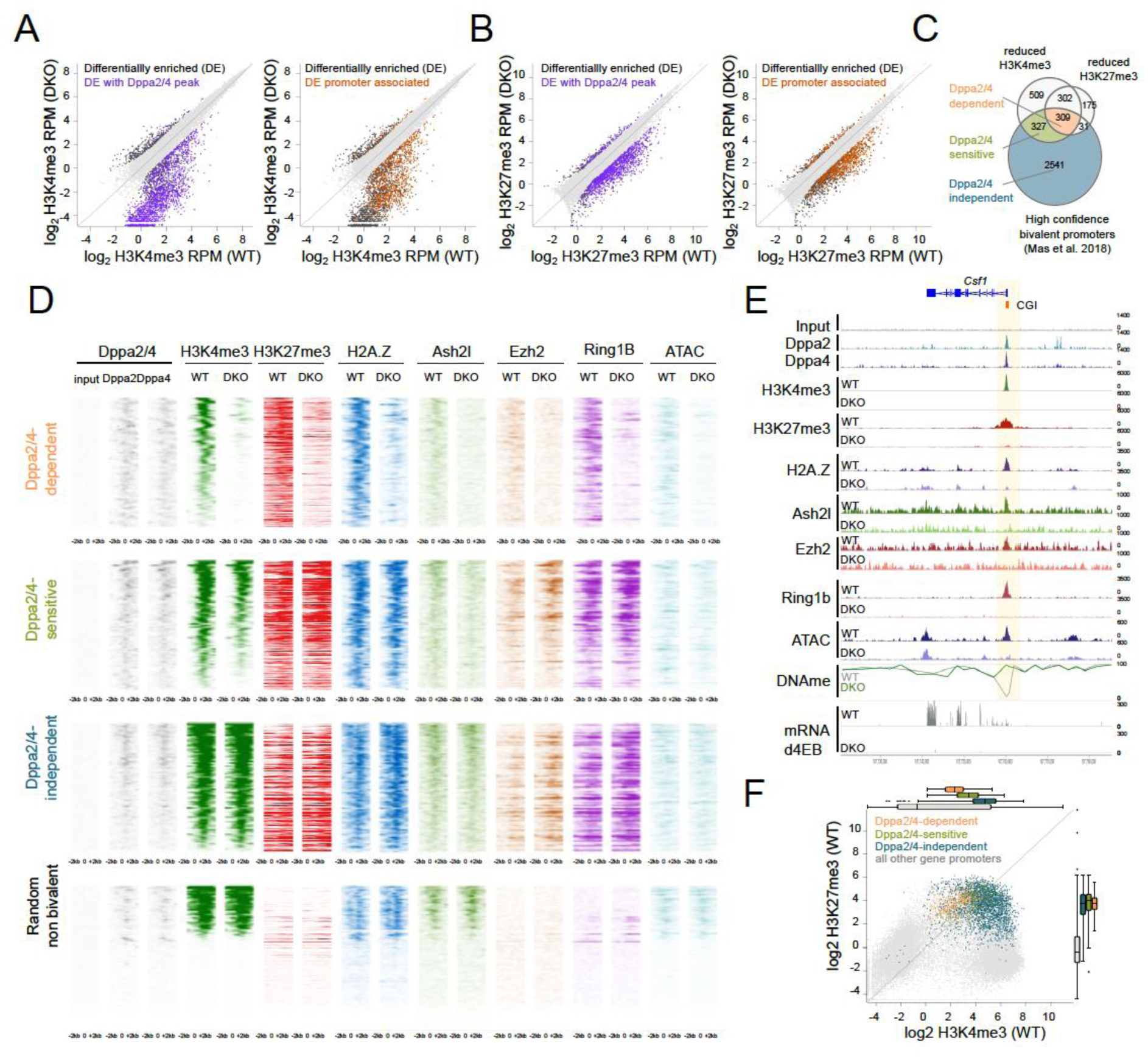
Dppa2/4 are required to maintain bivalent chromatin at a subset of developmental genes. (A) Scatter plots showing H3K4me3 enrichment as log_2_ RPM at H3K4me3 peaks in WT and Dppa2/4 DKO cells. Differentially enriched peaks (DESeq2) with >2-fold change are shown in dark grey with those overlapping Dppa2/4 peaks (purple) or promoters (orange) highlighted. (B) similar to A but showing H3K27me3 peaks. (C) Overlap between promoters (TSS +/-1kb) with reduced H3K4me3 or H3K27me3 enrichment in Dppa2/4 DKO cells (this study) with bivalent gene promoters defined by (*18*). (D) Aligned probe plots of wild type (WT) and Dppa2/4 double knockout (DKO) ESCs showing Dppa2 and Dppa4 (grey), H3K4me3 (dark green), H3K27me3 (dark red), H2A.Z (blue), Ash2L (light green), Ezh2 (orange) and Ring1b (purple) enrichment start of gene +/-2kb at Dppa2/4-dependent (top) and Dppa2/4-independent (middle) promoters versus a random subset of not bivalent promoters (bottom). Chromatin accessibility measured by ATACseq is shown in light blue. All data from this study. (E) Genome browser view of a Dppa2/4-dependent gene locus *Csf1* showing different chromatin marks, chromatin accessibility (ATAC-seq), DNA methylation (DNAme) and transcription at day 4 of embryoid body differentiation are shown for wild type (WT) and Dppa2/4 double knockout (DKO) ESCs. Promoter region is highlighted in pale yellow, CpG island (CGI) denoted by orange box. (F) Scatterplot showing levels of H3K4me3 and H3K27me3 in wild type cells at gene promoters, highlighting Dppa2/4-dependent (apricot), -sensitive (green) and -independent (teal) promoters. Box plots of are shown on top (H3K4me3) and right (H3K27me3).

We termed those bivalent promoters that had lost both H3K4me3 and H3K27me3 “Dppa2/4-dependent”, and those that had lost just H3K4me3 “Dppa2/4-sensitive” to distinguish them from “Dppa2/4-independent” bivalent promoters which remained unaltered (Fig. 3C-D). Consistent with their epigenetic changes, Dppa2/4-dependent promoters failed to recruit COMPASS member Ash2l, and the Polycomb members Ring1B and Ezh2, while Dppa2/4-sensitive promoters had reduced Ash2l levels but unaltered Ring1B and Ezh2 levels (Fig. 3D-E, S3B-F). Chromatin accessibility was substantially reduced at Dppa2/4-dependent promoters and partially impaired at Dppa2/4-sensitive promoters, whilst the remaining bivalent promoters remained largely unaffected (Fig. 3D-E, S3C, S3E-G). Consistent with the association between Dppa2/4 and the SRCAP complex, Dppa2/4-dependent promoters also exhibited lower H2A.Z enrichment (Fig. 3D-E, S3B-F). Interestingly, the response of a gene to Dppa2/4 loss was correlated with the levels of H3K4me3 in WT cells, with Dppa2/4-dependent genes having the lowest, Dppa2/4-sensitive genes intermediate and Dppa2/4-independent genes moderate-to-high H3K4me3 levels. H3K27me3 levels were not indicative of the gene’s response (Fig. 3D, F). Together, these findings support a role for Dppa2/4 in recruiting and/or stabilising Polycomb and COMPASS complexes at a subset of promoters to maintain a bivalent chromatin structure.

### Dppa2/4-dependent bivalent genes are characterised by low H3K4me3, low expression and initiating but not elongating RNA polymerase II

Next, we sought to understand what distinguished Dppa2/4-dependent from -sensitive and - independent bivalent genes. We took an unbiased approach and used machine learning algorithms to develop models that could predict based on genomic and epigenomic features (Table S5) the epigenetic effect of losing Dppa2/4. The initial 3-class Random Forest classification model had an overall accuracy of 72% and was able to classify Dppa2/4-dependent and Dppa2/4-independent genes. However Dppa2/4-sensitive genes were frequently misclassified, likely due to their high similarity to Dppa2/4-dependent promoters (Fig. 4A). We therefore focused on the two extreme categories (Dppa2/4-dependent and -independent), generating a 2-class Random Forest Classification model with 90% accuracy (Fig. 4B).

**Figure 4:**
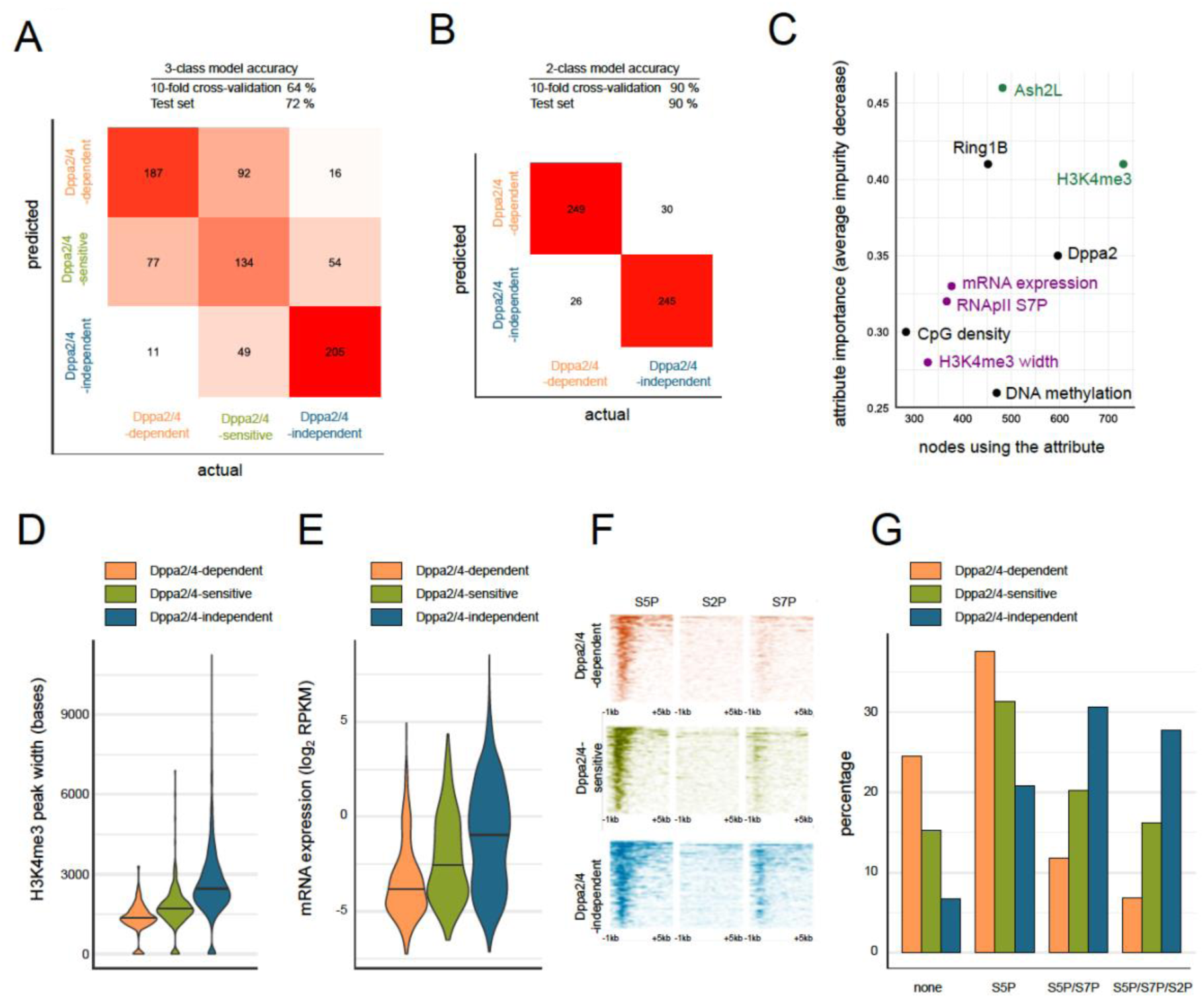
Dppa2/4-dependent bivalent genes characterised by low H3K4me3, low expression and initiating but not elongating RNA polymerase II. (A, B) Overall accuracy and confusion matrices for Random Forest promoter classification predicting either three (A) or two classes (B). The heatmap shows numbers of correctly and incorrectly classified promoters from a class balanced training set. (C) Ranking of the most predictive attributes in the 2-class Random Forest model showing average impurity decrease and number of nodes using each attribute. (D) H3K4me3 peak width in wild type ESCs. (E) Expression of genes in WT ESCs. (F) Aligned probe plots showing enrichment of different RNA polymerase II modifications at gene transcription start sites from 1kb upstream to 5kb downstream of TSS. Data reanalysed from (*24*). (G) Percentage genes with different combinations of RNApII modifications. Data reanalysed from (*24*).

Reassuringly, the top ranking attributes were directly related to levels of H3K4me3 (Fig. 4C), a feature we had already noticed was correlated with the response of a gene to loss of Dppa2/4 (Fig. 3D, F). Notably, the next group of attributes predictive of a gene’s response to loss of Dppa2/4 was directly linked to the transcriptional status of the gene (Fig 4C, Table S5). Dppa2/4-independent promoters had larger H3K4me3 peak width (Fig. 4D), a feature associated with increased transcriptional consistency yet independent of overall levels of gene expression (*23*) (Fig. S4A). Moreover, levels of gene expression (Fig. 4E) were higher for Dppa2/4-independent promoters than -dependent promoters. Also amongst the top storing attributes was phosphorylation of Serine 7 in the C-terminal domain (CTD) of RNApolymerase II (RNApII) (Fig. 4C). RNApII CTD phosphorylation status changes as it progresses through the transcriptional cycle: Serine 5 phosphorylation (S5P) correlates with initiating RNApII, while Serine 7 phosphorylation (S7P) is associated with active promoters and coding regions, and Serine 2 phosphorylation (S2P) with elongating RNApII (*24*). Strikingly, whilst both Dppa2/4-dependent and -independent promoters had initiating RNApII, only the Dppa2/4-independent promoters had progressed to actively elongating RNApII (Fig. 4F-G, S4B).

None of the other attributes, including Dppa2/4, Polycomb complexes or the Polycomb catalysed histone modifications H3K27me3 and H2AUb119 (Fig. S4C) showed large changes between Dppa2/4-dependent, -sensitive and -independent genes. We also examined genetic features of Dppa2/4-dependent and -independent promoters. There were no large differences in repeat composition (Fig. S4D), nor were there any motifs consistently enriched in either group of promoters (data not shown). Consistent with their lower expression, Dppa2/4-dependent promoters had fewer CpG islands and a slight increase in CpG density and content, which corresponded to higher levels of DNA methylation (Fig. S4E), reflective of the transcriptional status of the genes. In summary, our analyses highlight the variable dependency of bivalent gene promoters on Dppa2/4, with those characterised by low levels of H3K4me3, initiating but not elongating RNApII and low expression losing their bivalent chromatin structure in the absence of Dppa2/4.

### Dppa2/4-dependent bivalent promoters gain DNA methylation and can no longer be activated upon differentiation

The bivalent genes affected by loss of Dppa2/4 include many important developmental genes. Of those 310 Dppa2/4-dependent genes for which knockout mice have been generated and characterised, 65 (21%) have either complete or partial lethality, often at postnatal/preweaning stages. 15 (5%) of those are associated with lung defects and 31 (10%) with skeletal defects (www.informatics.jax.org). Therefore, the precise temporal regulation of these genes is crucial for proper development. To probe the relationship of bivalency in early development and subsequent functional readout, we analysed the functional and mechanistic consequences of altering chromatin bivalency at Dppa2/4-dependent and -sensitive genes during embryoid body differentiation. Normally, in WT cells, Dppa2/4-dependent, -sensitive and -independent bivalent promoters are upregulated over 9 days (Fig. 5A). However, in Dppa2/4 DKO cells, Dppa2/4-sensitive genes fail to be efficiently upregulated whilst the Dppa2/4-dependent bivalent genes are not transcribed and remain silent (Fig. 5A). To understand why these genes are repressed in the absence of Dppa2/4, we profiled DNA methylation, an epigenetic layer associated with promoter silencing. Global DNA methylation levels were similar between WT and DKO cells (Fig. 5B, S5A), however we observed that a subset of regions enriched for promoters and gene bodies gained DNA methylation in the DKO cells (Fig. 5B-C). Remarkably, these regions corresponded to Dppa2/4-dependent and -sensitive promoters which gained high and intermediate levels of DNA methylation in Dppa2/4 DKO cells respectively, whereas Dppa2/4-independent promoters were unaffected (Fig. 5D). Consistently, previous work revealed increased promoter DNA methylation and H3K9me3 at 4 promoters, including the Dppa2/4-dependent gene Nkx2-5, in ESCs lacking Dppa2 (*13*). This likely explains why Dppa2/4-dependent promoters can no longer be activated upon differentiation.

**Figure 5:**
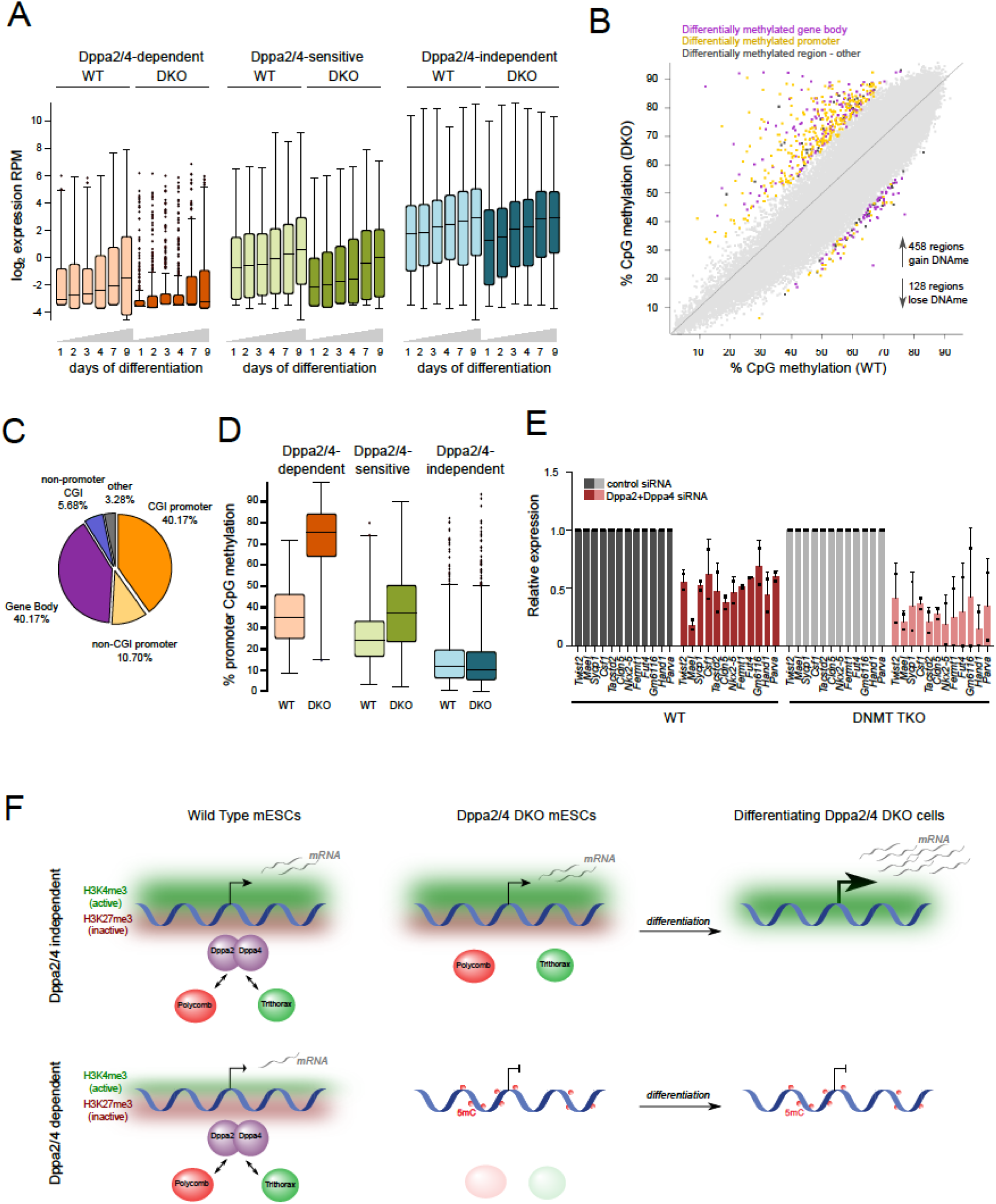
Dppa2/4-dependent promoters gain DNA methylation and fail to be upregulated during differentiation. (A) Expression of Dppa2/4-dependent (apricot), -sensitive (green) and -independent (blue) genes between WT (light) and Dppa2/4 DKO (dark) cells during 9 days of embryoid body differentiation. (B) Scatterplot showing DNA methylation levels in 100 CpG running windows between WT and Dppa2/4 DKO cells highlighting differentially methylated gene bodies (purple), promoters (yellow) and other regions (dark grey). (C) Genome features associated with hypermethylated probes. (D) DNA methylation levels at Dppa2/4-dependent (apricot), -sensitive (green) and -independent (blue) gene promoters between WT (light) and Dppa2/4 DKO (dark) cells. (E) Expression of Dppa2/4-dependent genes following control (grey) or Dppa2/4 (red) siRNA treatment in WT (left, dark) and Dnmt TKO (right, light) cells. Relative expression is normalised to the level of control siRNA (dark bars) for WT and DNMT TKO cells. Dots represent biological replicates and bars averages plus standard deviation. (F) Summary of epigenetic features and transcriptional consequences of Dppa2/4-dependent and Dppa2/4-independent bivalent promoters.

Finally, we sought to understand whether Dppa2/4-dependent genes first lost bivalency then gained DNA methylation or *vice versa*. We separated the two molecular events by knocking down Dppa2/4 in ESCs lacking the three DNA methyltransferases (DNMT TKO cells) and consequently have no detectable 5-methylcytosine or 5-hydroxymethylcytosine (*25*) (Fig. S5B-C). Remarkably, whilst similarly expressed in untreated WT and DNMT TKO cells (Fig. S5D), siRNA knockdown of Dppa2/4 (Fig. S5B, E) led to a downregulation of Dppa2/4-dependent genes in both WT and DNMT TKO cells (Fig. 5E), revealing that DNA methylation is not required for their downregulation. This suggests that DNA methylation is a consequence, not a cause of bivalency loss at these regions.

## Conclusions

It is currently unclear how both H3K4me3 and H3K27me3 modifications are targeted to bivalent chromatin in pluripotent cells, and the functional significance of this for development. We reveal that the heterodimerising proteins Dppa2 and Dppa4 function as epigenetic priming factors by regulating chromatin bivalency at over 600 bivalent promoters in ESCs. Regions that lose the bivalent H3K4me3 and H3K27me3 modifications subsequently gain repressive DNA methylation and fail to become active upon differentiation (Fig. 5F). Importantly, our study reveals a targeting principle for bivalent chromatin to a set of important developmental gene promoters, and the signficiance losing this structure has on cell fate acquisition.

In contrast to the stereotypical hetereogeneous nature of serum/LIF grown ESCs (*14–16*), ESCs lacking Dppa2/4 are transcriptionally more homogeneous and have an impaired ability to efficiently differentiate. Bivalent genes are inherently more dynamic and transcriptionally heterogeneous in ESCs (*26*), and it will be exciting to decipher the extent to which Dppa2/4 contribute to this. Our results also raise the possibility that bivalent promoters differ in their stability, with Dppa2/4-dependent promoters being potentially less robust, consistent with the suggestion that bivalent chromatin operates as a bistable system switching rapidly between active and silent states (*27*).

Importantly, Dppa2/4 target both H3K4me3 and H3K27me3 and interact with the catalytic machinery for both these modifications. Interactions between Dppa2/4 and Polycomb/COMPASS are likely transient and sub-stoichiometric implying that Dppa2/4 may act by recruiting and/or stabilising these complexes to their target loci, rather than existing as stable members of these complexes. Furthermore, Dppa2/4 also directly interacts with the SRCAP complex that deposits H2A.Z and has also been shown to associate with MLL and Polycomb complexes (*28–31*), suggesting another mechanism by which Dppa2/4 may target the bivalent chromatin machinery in ESCs.

Intriguingly, bivalent genes have a range of responses to loss of Dppa2/4 in ESCs. Through machine learning approaches we reveal that Dppa2/4-dependent bivalent promoters that lose both H3K4me3 and H3K27me3 are characterised by lower H3K4me3 enrichment and breadth, reduced gene expression and absence of elongating RNA polymerase II in WT cells. We hypothesise that these transcriptionally inert promoters may require continuous targeting of Polycomb and COMPASS machinery by Dppa2/4 to maintain the primed bivalent chromatin state. In contrast, positive feedback loops at actively transcribed Dppa2/4-independent genes may reinforce their bivalent epigenetic structure, rendering them immune to loss of Dppa2/4. The slightly higher expression and H3K4me3 levels at Dppa2/4-sensitive bivalent promoters which lose just H3K4me3 likely explains why these are not as affected as Dppa2/4-dependent promoters in Dppa2/4 DKO cells.

Our study identifies Dppa2/4 as epigenetic priming factors which function by establishing a permissive epigenetic landscape in pluripotent cells enabling appropriate activation of gene expression programmes at future stages of development. Dppa2/4 DKO ESCs fail to efficiently differentiate, likely due to the loss of bivalent chromatin and gain in DNA methylation at important developmental promoters. Consistently, zygotic knockouts for Dppa2/4 survive embryogenesis only to succumb shortly after birth from defects in tissues in which these proteins are not expressed (*12, 13*), although maternal stores of Dppa2/4 may be masking more severe developmental defects in gastrulating embryos. Dppa2/4 have additional roles in regulating the zygotic transcriptional programme *in vitro* (*6–8*), and enhance iPSC reprogramming (*9*) suggesting their roles as epigenetic priming factors may extend more generally to processes that encompass cell fate transitions. It will be exciting to determine how many other proteins, similar to Dppa2/4, act as epigenetic priming factors, establishing a permissive chromatin landscape at important developmental genes in pluripotent cells, enabling their effective and timely activation at later temporal stages.

## Acknowledgements

We thank all members of the Reik laboratory for helpful discussions. We also thank Federico di Tullio for help generating overexpression cell lines, Felix Krueger for processing sequencing data and general bioformatics support, Steven Wingett for bioinformatic assistance and Simon Andrews for bioinformatic advice. We thank Bethan Hussey and Elizabeth Easthope at Sanger Institute and Kristina Tabbada at Babraham Institute for assistance with high-throughput sequencing, Rachael Walker for assistance with flow cytometry, Judith Webster and David Oxley for mass spectrometry. DNMT TKO cells were a kind gift from Dirk Schübeler (FMI). M.E.-M. was supported by an EMBO Fellowship (ALTF938-2014) and a Marie Sklodowska-Curie Individual Fellowship. A.P. is supported by Sir Henry Wellcome Fellowship (215912/Z/19/Z). M.B. was supported by an Erasmus Grant. Research in the Reik laboratory is supported by the Biotechnology and Biological Sciences Research Council (BB/K010867/1), Wellcome Trust (095645/Z/11/Z), and European Union Epigenesys.

## Author contributions

M.A.E.-M. and W.R. conceived, designed and supervised the study. M.A.E.-M. performed experiments, analysed data and wrote the paper. A.P. helped perform ChIP-qPCR, ChIP-seq, Co-IP and Western blotting experiments, prepared cells for proteomics and analysed proteomic and ChIP-seq data. M.B. performed embryoid body differentiation assays and qPCRs. C.K. analysed chomatin state association of Dppa2/4, helped with scRNA-seq analysis, and generated the Random Forest Model for Dppa2/4 dependency based on promoter features. V.N.R.F. and C.D’S. performed qPLEX-RIME experiments. S.C. generated scNMT-seq libraries.

Sequencing data has been deposited in GEO and proteomics data under PRIDE identifier PXD014981.

W.R. is a consultant and shareholder of Cambridge Epigenetix. The remaining authors declare no competing financial interests.

## SUPPLEMENTAL MATERIALS

### Materials and Methods

#### Cell culture and flow cytometry

E14 mouse ESCs were grown using standard serum/LIF culture conditions (DMEM, 4,500 mg/L glucose, 4 mM L-glutamine, 110 mg/L sodium pyruvate, 15% fetal bovine serum, 1 U/mL penicillin, 1 mg/mL streptomycin, 0.1 mM nonessential amino acids, 50 mM b-mercaptoethanol, 10^3^ U/mL LIF) at 37 degrees Celsius in normal oxygen, feeder-free on gelatin coated plates. Stable overexpression cell lines were generated by transfecting GFP, Dppa2-GFP or Dppa4-GFP constructs previously described (*6*) using Lipofectamine 2000 on preplated cells and resistant cells selected with appropriate antibiotics for at least 1 week. Resistant cells were sorted as single cells into 96 well plates by flow cytometry using a BD Aria III or BD Influx high-speed cell sorter, and clonal expansion. Overexpression was validated by qPCR and Western Blotting. siRNA transfections were performed by transfecting Dharmacon siRNA ON-TARGETplus siRNA SMARTpool at a final concentration of 50 nM with Lipofectamine. CRISPR double knockout ESC line (clone 43) was described previously (*6*), and additional CRISPR double knockout ESC lines (clone 37 and clone 53) generated as previously described (*6*). WT and DNMT TKO cell lines were described in (*25*). For embryoid bodies, 2×10^6^ mESCs were cultured on 10cm low-attachment dishes in standard ESC medium containing all described components except LIF.

#### RNA isolation, qPCR and RNA-seq

Total RNA was isolated using TriReagent (Sigma) or RNA-DNA allprep columns (Qiagen) 0.5-1μg DNAse treated (Thermo Fisher EN0525) RNA was converted to cDNA using random priming (Thermo RevertAid K1622). qRT-PCR was performed using Brilliant II or II SYBR master mix (Agilent Technologies) and relative quantification performed using the comparative CT method with normalisation to CycloB1 levels. Primer sequences available upon request. Opposite strand-specific polyA RNA libraries were made using 1 µg of DNase-treated RNA at the Sanger Institute Illumina bespoke pipeline and sequenced using the Illumina HiSeq2 platform. EB RNA-seq raw FastQ data were trimmed with Trim Galore (version 0.4.4, default parameters) and mapped to the mouse GRCm38 genome assembly using Hisat2 version 2.1.0. Stable overexpression clone data used Trim Galore v0.5.0_dev and HiSat2 v2.1.0.

#### Single-cell RNA-sequencing library preparation and analysis

Single cells were sorted into individual wells of 96 well plates, ensuring both wild type and Dppa2/4 DKO cells were present on each plate to control for plate batch effects. RNA-sequencing libraries were prepared using scNMT-seq method as previously described (*34, 35*) and transcriptomes sequenced using the Illumina HiSeq platform to generate PE75bp reads. Reads were trimmed using Trim Galore version 0.5.0 using default parameters and aligned to the mouse GRCm38 genome using hisat2 version 2.1.0 using default parameters. Initial data analysis was performed using Seqmonk software. Poor quality cells with <20% genes covered, <90% exonic reads or >10% mitochondrial reads were excluded from analysis. A total of 148 wild type (69 from clone 57 and 79 from clone 58) and 57 Dppa2/4 DKO (all from clone 43) cells were used in downstream analysis. Total read counts ranged from 319,214 to 2,033,187 reads per cell with a total of 14,398 genes expressed (>1 raw read count in >10 cells). Cell-to-cell correlations were calculated in Seqmonk by calculating all pairwise Pearson correlation values between cells for expressed genes and determining the mean correlation value for each cell. For pseudotime analysis, raw read counts for scRNA-seq samples were exported from Seqmonk and analysed using the Monocle 3 package (*36–39*). Data was normalised using Principal Component Analysis (50 dimensions) and corrected for batch effects (Batcherlor) (*40*). Dimensionality reduction was performed using UMAP (*41*). Cells were ordered in pseudotime with the starting point of the trajectory (root node) selected near cells with high expression of naïve ES cell markers.

#### Chromatin immunoprecipitation

Ten million cells were fixed in 1% formaldehyde (Fisher Scientific 28906) in DMEM (Invitrogen 41966-052) for 10 minutes, quenched in 0.125M glycine, and scraped off cell culture dishes using cell scrapers. Cells were lysed with buffers LB1 (50mM Hepes-KOH (pH 7.5), 140mM NaCl, 1mM EDTA, 10% Glycerol, 0.5% Igepal CA-630, 0.25% Triton-X 100), LB2 (10mM Tris-HCl (pH 8.0), 200mM NaCl, 1mM EDTA, 0.5mM EGTA), and LB3 (10mM Tris-HCl (pH 8.0), 100mM NaCl, 1mM EDTA, 0.5mM EGTA, 0.1% Na-Deoxycholate, 0.5% N-lauroylsarcosine, 1x Protease Inhibitors (Roche)) consecutively, before fragmenting chromatin by sonicating for 30 cycles of 30s ON high /30s OFF (Diagenode Bioruptor). For each ChIP, 5µg antibody (anti-Ash2l Bethyl Labs A300-489, anti-Ezh2 CST D2C9, anti-H2A.Z Abcam ab4174, anti-H3K4me3 Abcam ab8580, anti-H3K27me3 Active Motif AM39155, anti-IgG Abcam ab125938, anti-Ring1b CST D22F2) was pre-bound to protein G dynabeads (Invitrogen) and blocked with 0.5% BSA in PBS. Sonicated DNA was added to antibody-bound beads with 10% Triton-X 100 and incubated overnight at 4°C on a rotator. Beads and DNA were washed 6 times with RIPA buffer (50mM HEPES-KOH (pH 7.6), 1mM EDTA, 0.7% Na-Deoxycholate, 1% Igepal CA-630, 0.5M LiCl), followed by one wash in 1x TBS and overnight incubation at 65°C with elution buffer (50 mM Tris-HCl, pH 8; 10 mM EDTA; 1% SDS). For precipitation of chromatin, samples were treated with RNase A and Proteinase K. DNA was purified using MinElute PCR purification columns (Qiagen 28006) and eluted in 30µL of TE. DNA was quantified using HS DNA Qubit (ThermoFisher) or PicoGreen (ThermoFisher) assays and analysed using qPCR (*Twist2-F* GGAGCGGTTGTCAAAACGTC, *Twist2-R* CTTGAACGCCCTAGCATCCA, *Fermt1-F* AGCGGGTCCAGTGATGTTG*, Fermt1-R* CCTTCTCCTACTCGGAGCGA, *Klf4-F* GAAAGTCCTGCCACGGGAA, *Klf4-R* CTGGATGAGTCACGCGGATAA) or libraries generated using MicroPlex Library Preparation Kit (Diagenode) following the manufacturer’s instructions. Libraries were quality controlled using bioanalyser HS DNA chips (Agilent) and single-end 50bp reads sequenced using the Illumina HiSeq2 sequencing platform. Raw FastQ data were trimmed with Trim Galore v0.6.1 and aligned to mouse GRCm38 genome using Bowtie 2 v 2.3.2.

#### ATAC-seq

ATAC-seq libraries were performed as previously described (*42*) with the following modification to use on 10,000 cells: initial transposition reaction was performed with 10,000 cells, 10μl 2x TD buffer, 0.5μl Tn5 enzyme and 9.5μl H_2_O for 30minutes at 37 degrees Celsius. A total of 15 cycles of amplification were used. Two technical replicates were performed per clone, with 3 WT and 3 Dppa2/4 DKO clones used (total of 12 samples). 75bp paired-end reads were sequenced using the Illumina HiSeq2 platform. Raw FastQ data were trimmed with Trim Galore (v0.5.0_dev) using standard parameters and aligned using Bowtie 2 v2.3.2. Data was analysed using Seqmonk, graphing and statistics was performed using Excel, RStudio or Graphpad Prism8.

#### DNA methylation analyses

Genomic DNA quantified using picogreen assay (Invitrogen) was digested using DNA Degradase plus (Zymo Research) overnight at 37 degrees and analysed by liquid chromatography-tandem mass spectrometry on a LTQ Orbitrap Velos mass spectrometer (Thermo Scientific) fitted with a nanoelectrospray ion-source (Proxeon, Odense, Denmark). Mass spectral data for cytosine, 5-methylcytosine and 5-hydroxymethylcytosine were acquired as previously described (*43*). Whole genome bisulfite libraries were generated using NOME-seq kit from Active Motif (103946) according to manufacturer’s instructions with 6 cycles of amplification for 2 WT clones (clones 57 and 58) and 1 Dppa2/4 DKO clone (clone 43). Unfortunately, the GC methyltransferase reaction did not work adequately so only DNA methylation (and not accessibility) information was analysed. Raw fastQ data were trimmed using Trim Galore version 0.4.4 using standard parameters, reads aligned using Bismark v0.18.2 (*44*) and data analysed using Seqmonk software.

#### Western Blotting

Protein lysates were generated by resuspending cells in Laemmli buffer (62.5 mM Tris-HCl pH 6.8, 2% SDS, 5% β-mercaptoethanol, 10% glycerol) without dye, boiling for 3-5 minutes at 98°C and immediately storing at −80°C. Protein quantification was performed using a Bradford Assay (Bio-Rad). Samples were boiled prior to loading onto NuPAGE Bis-Tris gels (Invitrogen NP0322BOX) together with NuPAGE MES SDS (Invitrogen NP0002) or MOPS SDS running buffer (Invitrogen NP0001) and blotted on PVDF membranes. Following blocking in 5% skimmed milk/0.01%Tween/PBS, membranes were incubated with primary antibodies overnight (anti-β-Actin Abcam ab6276 1:20,000; anti-Ash2l Bethyl A300-489A 1:2,000; anti-Dmap CST 13326 1:1,000; anti-Dppa2 R&D MAB4356 1:500; anti-Dppa4 Santa Cruz sc-74614 1:200; anti-Ezh2 CST D2C9 1:1,000; anti-H2A.Z Abcam ab4174 1:1,000; anti-H3 Abcam ab1791 1:1000; anti-H3K4me3 Diagenode C15410003 1:1,000; anti-H3K27me3 Millipore 07-449 1:1000; anti-Hsp90 Abcam ab13492 1:5,000; anti-Mll2 CST D6X2E #63735 1:1000, anti-Ruvbl1 Abcam ab51500 1:100; anti-Suz12 CST D39F6 1:1,000). After extensive washing in PBS/0.01% Tween, horseradish peroxidase-conjugated secondary antibodies (anti-mouse – Santa Cruz sc-2005, 1:5000; anti-rabbit – Biorad 170-6515, 1:5000) were added for 1 h at room temperature, before further washing and detection, which was carried out with enhanced chemiluminescence (ECL) reagent (GE Healthcare, RPN2209).

#### Co-immunoprecipitation

Cells were washed extensively in cold PBS and collected by adding 500µL of Co-IP Lysis Buffer A (10mM Hepes-KOH, pH7.9, 1.5mM MgCl2, 10mM KCl, 0.5mM DTT, 0.05% NP-40, 250u/µL Benzonaze +_C0mplete protease inhibitors (Roche)) directly to a 15cm tissue culture plate before scraping and transferring to falcon tubes. After incubating on ice for 10 minutes, lysed cells were collected by centrifugation at 3,000rpm for 10 minutes at 4°C. The pellet was suspended in 374µL of Co-IP Lysis Buffer B (5mM Hepes-KOH, pH7.9, 1.5mM MgCl2, 0.2mM EDTA, 0.5mM DTT, 26% glycerol (v/v), 250u/µL Benzonase +C0mplete protease inhibitors (Roche) and a further 26µL of 4.6M NaCl was added to lyse nuclear membranes. The resulting nuclear lysates were dounce homogonised (20 strokes) to further lyse membranes and shear DNA before a further 30-minute incubation on ice. Lysates were centrifuged at 24,000g for 20 minutes at 4°C and the supernatant collected and protein concentration quantified by Bradford assay (Bio-Rad). For immunoprecipitation, 50µL of protein G dynabeads (Invitrogen) were washed in IP wash buffer (10mM Tris-HCl pH7.5, 150mM NaCl, 0.5mM EDTA) and 5µg of anti-Dppa4 antibody (R&D AF3730) added before incubating at 4°C for 1 hour. Beads were washed 3 times in IP wash buffer to remove unbound antibody and 250µg of nuclear lysate was added, before further incubation at 4°C for 1 hour. Beads were washed 5 times in IP wash buffer to remove unbound protein before bound proteins were eluted from the beads by adding 50µL of 1x Laemmli buffer and boiling at 70°C for 10 minutes. Inputs were prepared by taking 25µg of nuclear lysate and adding an appropriate volume of 5x Laemmli buffer. Samples and inputs were analysed by western blotting as described above.

#### qPLEX-RIME

For qPLEX-RIME experiments 50 million cells were fixed, lysed and immunoprecipitated as described for *Chromatin immunoprecipitation* (see above). Following immunoprecipitation using 5µg of anti-GFP antibody (Abcam ab290), beads were washed 10 times in RIPA buffer and 2 times in 100mM ammonium bicarbonate (AMBIC) solution. Samples were prepared as described previously (*21*). In brief, after on bead tryptic digestion, C18 cleaned peptides were labelled with the TMT-10plex reagents (Thermo Scientific) for 1 hour. Samples were mixed and fractionated with Reversed-Phase cartridges at high pH (Pierce #84868). Nine fractions were collected using different elution solutions in the range of 5–50% ACN.

Peptide fractions were reconstituted in 0.1% formic acid and analysed on a Dionex Ultimate 3000 UHPLC system coupled with the nano-ESI Fusion Lumos mass spectrometer (Thermo Fisher Scientific, San Jose, CA). Samples were loaded on the Acclaim PepMap 100, 100 μm × 2 cm C18, 5 μm, 100 Å trapping column with the ulPickUp injection method using the loading pump at 5 μL/min flow rate for 10 min. For the peptide separation the EASY-Spray analytical column 75 μm × 25 cm, C18, 2 μm, 100 Ȧ column was used for multi-step gradient elution at a flow rate of 300 nl/min. Mobile phase (A) was composed of 2% acetonitrile, 0.1% formic acid and mobile phase (B) was composed of 80% acetonitrile, 0.1% formic acid. Peptides were eluted using a gradient as follows: 0 - 10 min, 5 % mobile phase B; 10 – 90 min, 5 – 38% mobile phase B; 90 −100 min, 38% - 95% B; 100 - 105 min, 95% B; 105 –110min, 95% - 5% B; 110 – 120 min, 5% B.

Data-dependent acquisition began with a MS survey scan in the Orbitrap (380 – 1500 m/z, resolution 120,000 FWHM, automatic gain control (AGC) target 3E5, maximum injection time 100 ms). MS2 analysis consisted of collision-induced dissociation (CID), quadrupole ion trap analysis, automatic gain control (AGC) target 1E4, NCE (normalized collision energy) 35, q-value 0.25, maximum injection time 50 ms, an isolation window at 0.7, and a dynamic exclusion duration of 45s. MS2–MS3 was conducted using sequential precursor selection (SPS) methodology with the top10 setting. HCD-MS3 analysis with MS2 isolation window 2.0 Th. The HCD collision energy was set at 65% and the detection was performed with Orbitrap resolution 50,000 FWHM and in the scan range 100–400 m/z. AGC target 1E5, with the maximum injection time of 105ms.

#### qPLEX-RIME data processing

The Proteome Discoverer 2.1. (Thermo Scientific) was used for the processing of CID tandem mass spectra. The SequestHT search engine was used and all the spectra searched against the Uniprot *Mus musculus* FASTA database (taxon ID 10090 - Version November 2018). All searches were performed using as a static modification TMT6plex (+229.163 Da) at any N-terminus and on lysines and Methylthio at Cysteines (+45.988Da). Methionine oxidation (+15.9949Da) and Deamidation on Asparagine and Glutamine (+0.984) were included as dynamic modifications. Mass spectra were searched using precursor ion tolerance 20 ppm and fragment ion tolerance 0.5 Da. For peptide confidence, 1% FDR was applied and peptides uniquely matched to a protein were used for quantification.

Data processing, normalisation and statistical analysis were carried out using qPLEXanalyzer (*21*) package from Bioconductor. Peptide intensities were normalised using median scaling and protein level quantification was obtained by the summation of the normalized peptide intensities. A statistical analysis of differentially-regulated proteins was carried out using the limma method (*45*). Multiple testing correction of p-values was applied using the Benjamini-Hochberg method (*46*) control the false discovery rate (FDR).

#### Next generation sequencing data processing and mapping

Aligned read (bam) files were imported into Seqmonk software (http://www.bioinformatics.babraham.ac.uk/ projects/seqmonk) for all downstream analysis using standard parameters. ChIPseq reads were extended by 250bp.

#### RNA-sequencing analysis

For RNA-seq analysis, data was quantified at the mRNA level using strand-specific quantification of mRNA probes using the RNA-seq quantification pipeline in Seqmonk. Differentially expressed genes were determined using both EdgeR (p-value of 0.05 with multiple testing correction) and intensity difference filter (p-value 0.05 with multiple testing correction), with the intersection between the two lists giving the high confidence differentially expressed genes.

#### ChIP-seq analysis

For ChIP-seq analysis, data was analysed using Seqmonk software. Peaks were called using MACS (p-value cutoff 1.0E-5, sonicated fragment size 300bp), and for Dppa2 and Dppa4 peaks, filtered further to retain peaks that had values above the 95% value of input (0.008) and a 2-fold change over input to retain 36 901 and 21 509 high confidence peaks for Dppa2 and Dppa4 respectively, or 39 218 peaks in total. For histone ChIP-seq analysis peaks were called using MACS (p-value cutoff 1.0E-5, sonicated fragment size 300bp) and differentially enriched peaks determined using DESeq2 (p-value 0.05, multiple testing correction). Unless otherwise specified, promoters were defined as the region covering 1kb up and 1kb downstream of the gene start or transcription start site (TSS).

#### Whole genome bisulfite sequencing analysis

Genome wide DNA methylation analysis was performed using 100 CpG running window probes and differentially methylated regions calculated using a binomial F/R filter with minimum 15% difference threshold, p-value <0.05. For promoter analysis, probes 1kb up and 1kb downstream of TSS were generated and methylation values calculated with a minimum count of 2 per position and at least 10 observations per feature, and combined using the mean.

#### Analysis of chromatin states

Published mouse ES cell chromatin states were used to annotate the GRCm38 genome (*47*). The states were generated using ChromHMM (*48*) and more information on the annotation can be found at https://github.com/guifengwei/ChromHMM_mESC_mm10/blob/master/README.md. Dppa2/4 peaks were annotated with the chromatin state that overlaps its centre. The percentage of Dppa2/4 peaks falling into each of the states was determined, and enrichment of the chromatin state over its genomic representation was calculated.

#### Machine learning on Dppa2/4 dependent and independent promoters

A Random Forest Classifier was trained to predict Dppa2/4 dependency on a class-balanced set of bivalent promoters (n=825 for 3-class prediction, n=550 for 2-class prediction). Attributes used are specified in Table S5. Attribute selection was performed using Correlation-based Feature Subset Selection, retaining 4/9 (3-class/2-class) out of 20 attributes. Performance was evaluated using 10-fold cross-validation and an independent test set (a random set of 10 % of all bivalent promoters which was not used for model generation).

Learning parameters: weka.classifiers.meta.AttributeSelectedClassifier -E ‘weka.attributeSelection.CfsSubsetEval -P 1 -E 1’ -S ‘weka.attributeSelection.BestFirst -D 0 -N 5’ -W weka.classifiers.trees.RandomForest ---P 100 -attribute-importance -I 100 -num-slots 1 -K 0 -M 1.0 -V 0.001 -S 1

#### Software

Data was analysed using Seqmonk software (http://www.bioinformatics.babraham.ac.uk/ projects/seqmonk), graphing and statistics was performed using Excel, RStudio (http://www.rstudio.com/), R (https://www.R-project.org/) or Graphpad Prism8. Weka (*49*) was used for machine learning.

#### Data availability

All sequencing data generated in this study has been submitted to GEO. The mass spectrometry proteomics data have been deposited to the ProteomeXchange Consortium via the PRIDE (*50*) partner repository with the data set identifier PXD014981. Dppa2 and Dppa4 ChIP data was reanalysed from (*9*), RNA polymerase II ChIP was reanalysed from (*24*), Ezh2 and Suz12 ChIP data in Figure 2 was reanalysed from (*32*), Ash2l-GFP ChIP data in Figure 2 reanalysed from (*1*), Mll2 ChIP data in Figure 2 reanalysed from (*2*), Tip60/Kat5 ChIP data reanalysed from (*33*), high confidence bivalent gene list from (*18*), DNMT TKO ESC RNAseq data from (*25*), 2C-like ZGA gene list from (*51*).

**Supplemental Figure 1, related to Figure 1.**
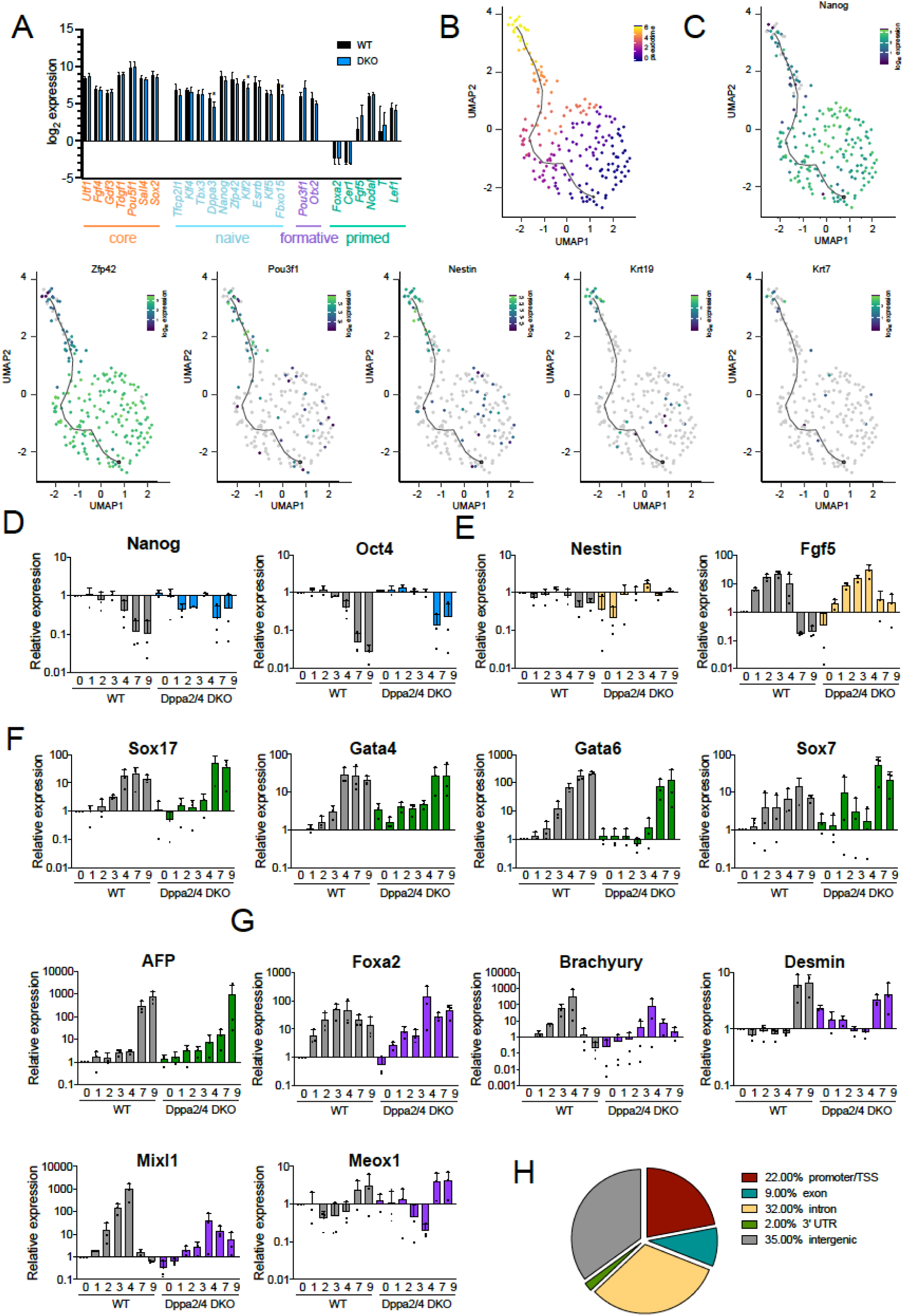
(A) Expression of core (orange), naïve (blue), formative (purple) and primed (aqua) pluripotency markers in WT (black) and Dppa2/4 DKO (blue) ESCs as determined by RNA-sequencing. 2-3 biological replicates are included for 3 individual clones for WT and DKO cells. * differences are statistically significant (DESeq2 + intensity difference filter with multiple testing correction) (B) Dimensionality reduction plot (UMAP) of single cell transcriptomes ordered by pseudotime (black line). (C) Expression of pluripotency and early differentiation markers in single cells ordered by pseudotime (black line) in UMAP dimensionality reduction plots. (D-G) Quantitative RT-PCR of pluripotency (D), ectoderm (E), endoderm (F) and mesoderm (G) markers during 9 day embryoid body differentiation between wild type (WT, grey) and Dppa2/4 DKO (coloured) cells. Error bars represent mean plus standard deviation of three independent differentiation experiments. RNA samples are the same that were used for RNA sequencing in Figure 1A-B. (H) Distribution of high confidence Dppa2/4 peaks. Data reanalysed from (*9*).

**Supplemental Figure 2, related to Figure 2.**
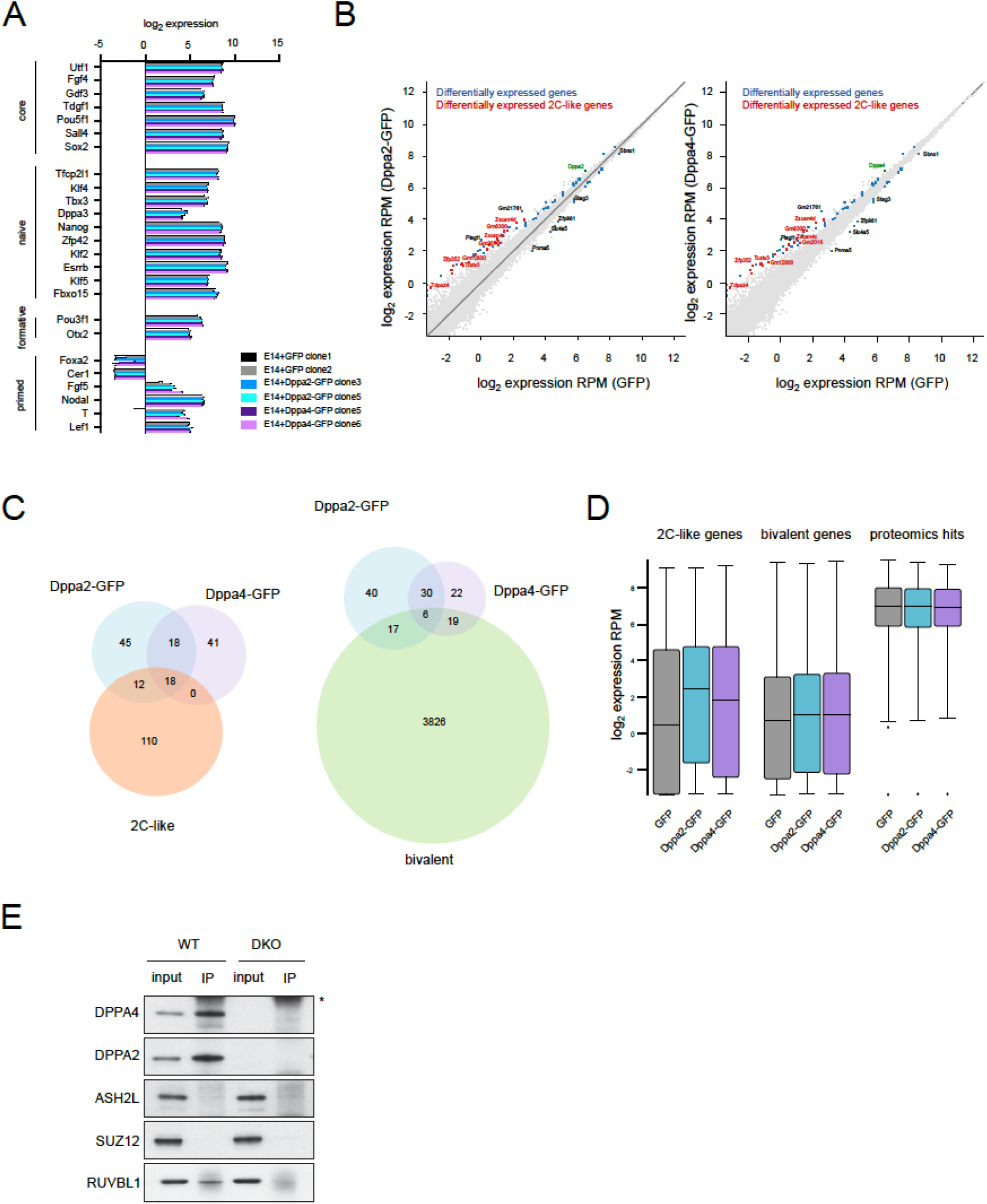
(A) Expression of core, naïve, formative and primed pluripotency markers in E14+GFP (black, grey), E14+Dppa2-GFP (blue) and E14-Dppa4-GFP (purple) overexpression clones as determined by RNA-sequencing. 2-3 biological replicates are included for each overexpression line. (B) Scatter plots showing gene expression between ESCs stably overexpressing GFP and Dppa2-GFP (left) or Dppa4-GFP (right). Differentially expressed genes are shown in blue, differentially expressed 2C-like ZGA genes are shown in red. Dppa2 and Dppa4 are shown in green. (C) Overlap in differentially expressed genes in Dppa2-GFP (blue) and Dppa4-GFP (purple) stable overexpression clones with 2C-like (left, orange) and bivalent (right, green) genes. (D) expression of 2C-like genes, bivalent genes and proteomics hits in GFP (grey), Dppa2-GFP (blue), Dppa4-GFP (purple) stable overexpressing cells. 2C-like genes from (*51*), bivalent genes from (*18*). Two independently targeted clones are shown per condition in all subpanels. (E) Endogenous Dppa4 immunoprecipitation (IP) followed by Western Blot in WT (first two columns) and Dppa2/4 DKO (last two columns) cells. * denotes IgG band.

**Supplemental Figure 3, related to Figure 3.**
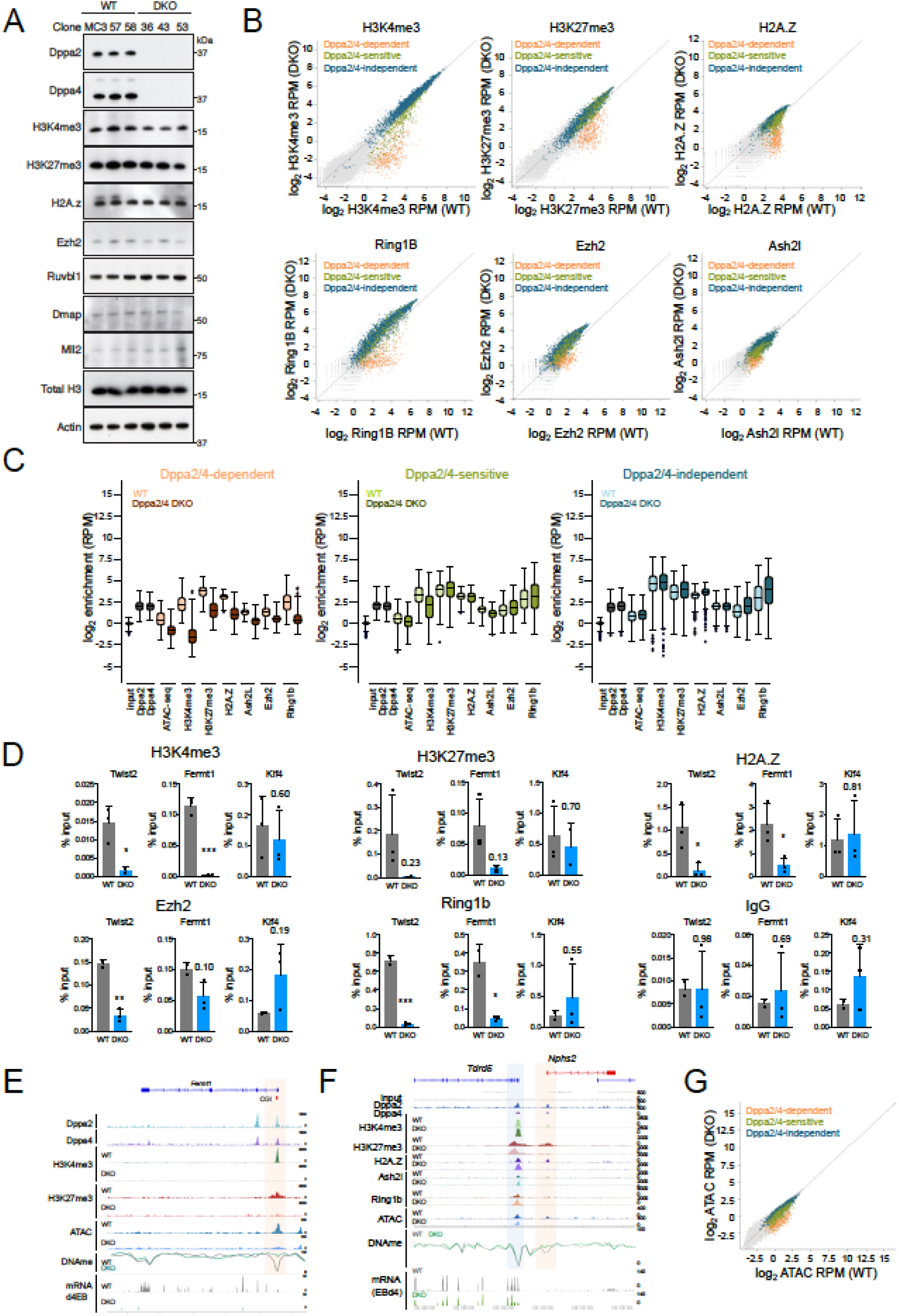
(A) Western blot showing protein levels of chromatin factors and histone modifications between three wild type (WT) ESC clones (left) and three Dppa2/4 DKO ESC clones (right). Histone H3 and β-actin are included as loading controls. (B) Scatter plots showing enrichment at gene promoters for H3K4me3 (top left), H3K27me3 (top middle), H2A.Z (top right), Ring1b (bottom left), Ezh2 (bottom middle) and Ash2l (bottom right) between WT (x-axis) and Dppa2/4 DKO (y-axis) ESCs. Dppa2/4-dependent (apricot), -sensitive (green) and -dependent (teal) genes are shown. (C) box-whisker plots showing enrichment of different chromatin proteins, histone modifications and chromatin accessibility measured by ATAC-seq at Dppa2/4-dependent (left, apricot), -sensitive (middle, green) and - independent (right, blue) promoters (TSS +/-1kb) comparing WT (light) and Dppa2/4 DKO (dark) ESCs. Input, Dppa2 and Dppa4 ChIP data from (*9*). All other data from this study. (D) ChIP-qPCR results at Dppa2/4-dependent genes Twist2 and Fermt1, compared to control Klf4 gene promoters in 2-3 biological replicates between WT (grey) and Dppa2/4 DKO (blue) cells for H3K4me3 (top left), H3K27me3 (top middle), H2A.Z (top right), Ezh2 (bottom left), Ring1B (bottom middle) and IgG control (bottom right). Error bars represent mean + standard deviation of 2-3 biological replicates. Individual data points are shown. * p<0.05, ** p<0.01, *** p<0.001 unpaired t-test. (E, F) Genome browser view of a Dppa2/4-dependent (Fermt1) (E), and a -dependent (Nphs2) and -independent (Tdrd5) gene (F). showing enrichment of chromatin modifications, proteins, chromatin accessibility (blue), DNA methylation (green/grey) and transcription (bottom two panels) are shown for wild type (WT) and Dppa2/4 double knockout (DKO) ESCs. (G) Accessibility measured by ATAC-seq at gene promoters (TSS +/-1kb) comparing Dppa2/4-dependent (apricot), -sensitive (green) and -independent (teal) promoters between WT (x-axis) and Dppa2/4 DKO (y-axis) cells.

**Supplemental Figure 4, related to Figure 4.**
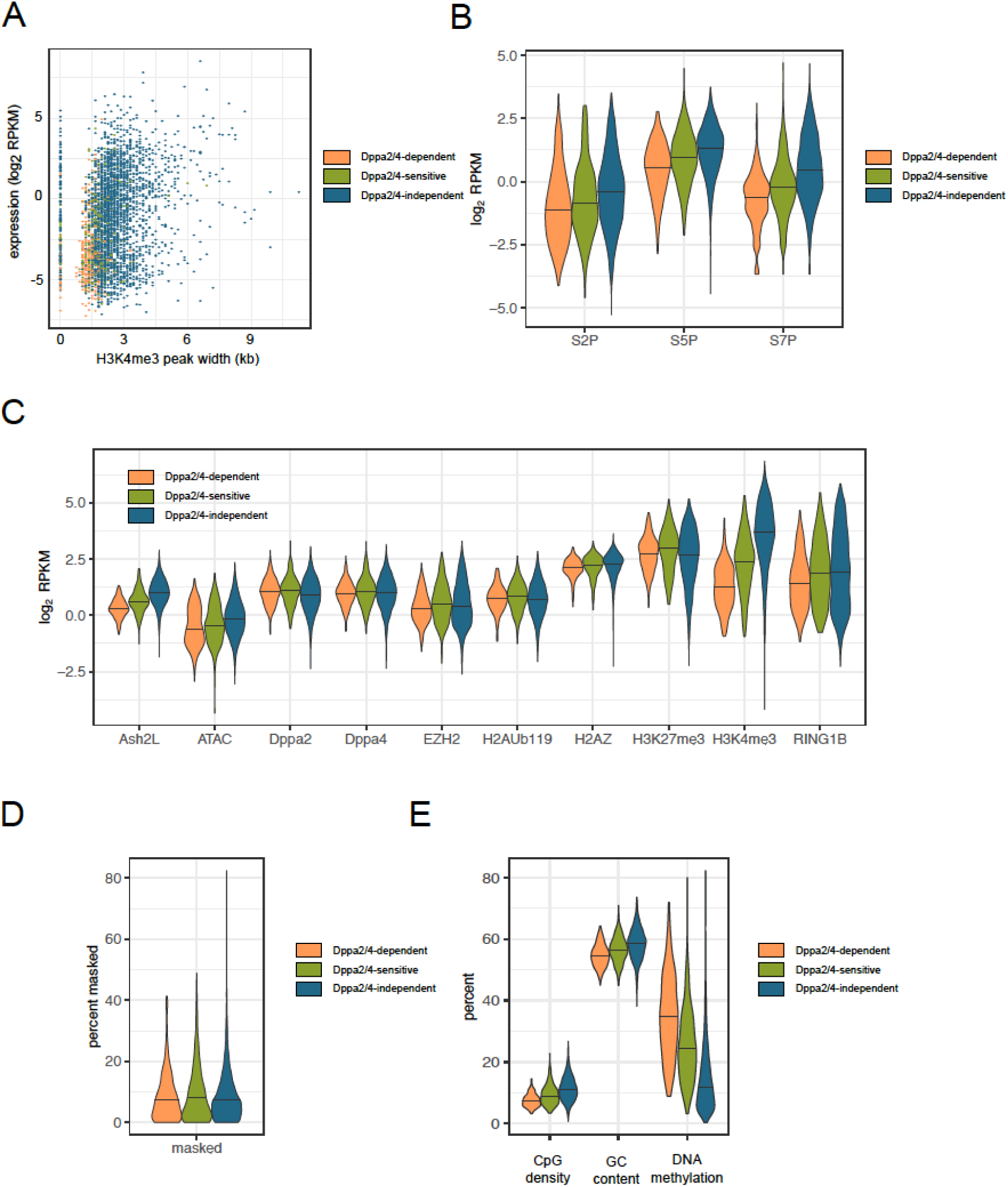
(A) Scatterplot comparing H3K4me3 peak width (x-axis) and expression of associated gene (y-axis) for Dppa2/4-dependent (orange), -sensitive (green) and -indpendent (blue) genes. (B) Enrichment for different RNA polymerase II modifications across gene bodies. Data reanalysed from (*24*). (C) Enrichment of different chromatin modifications and chromatin proteins along with chromatin accessibility (ATAC-seq) between Dppa2/4-dependent (orange), -sensitive (green) and -independent (blue) gene promoters (TSS +/-1kb). Dppa2/4 data reanalysed from (*9*), all other data from this study. (D) proportion of gene promoters (start of gene +/-1kb) sequence masked by repeats. (E) CpG density, GC content and DNA methylation levels at Dppa2/4-dependent (orange), -sensitive (green) and - independent (blue) gene promoters (TSS +/-1kb).

**Supplemental Figure 5, related to Figure 5.**
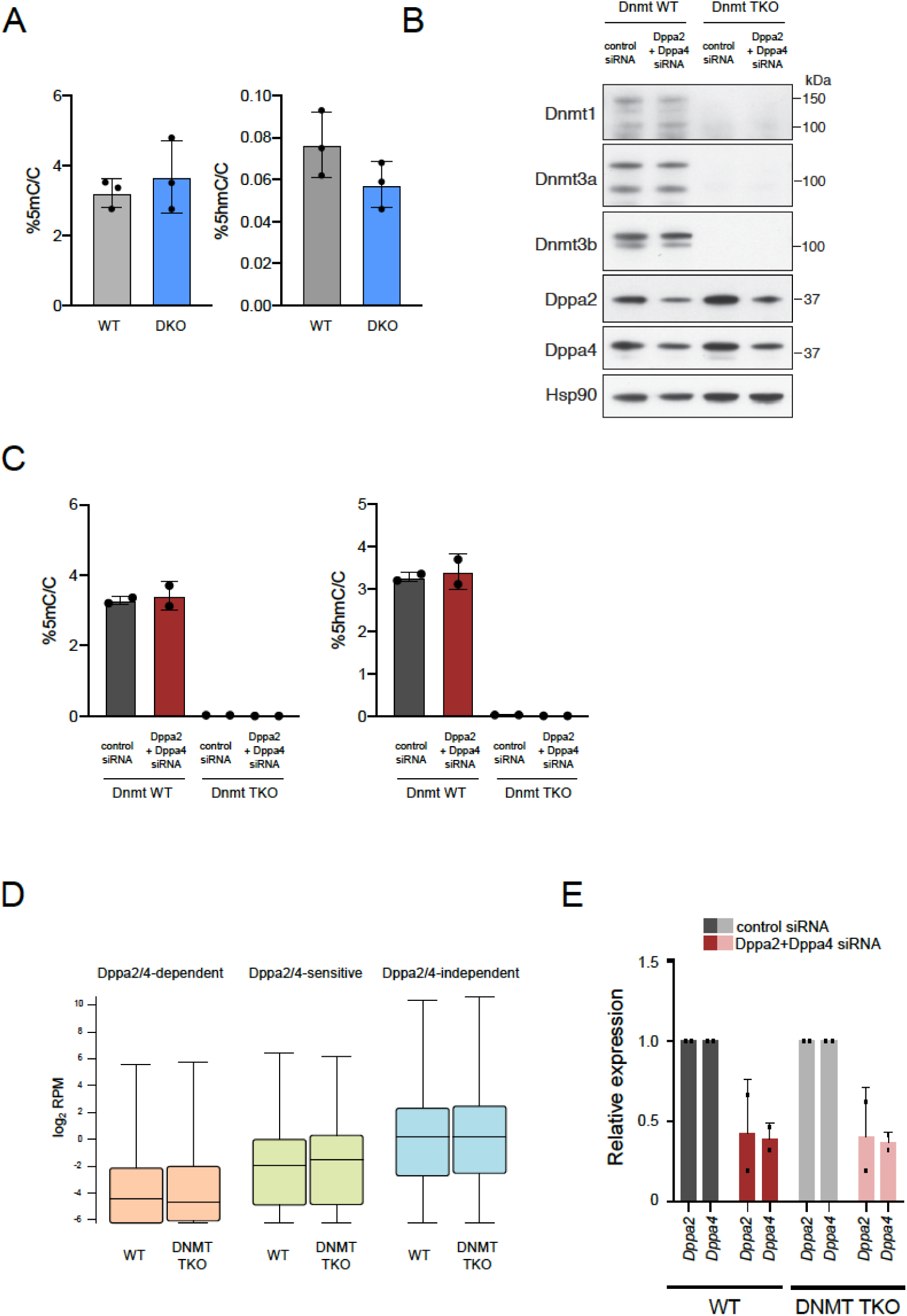
(A) Mass spectrometry quantification of 5-methylcytosine (5mC, left) and 5-hydroxymethylcytosine (5hmC, right) between WT (grey) and Dppa2/4 DKO (blue) cells. Bars represent mean +/-standard deviation of three independent clones. (B) Western blot showing expression of DNA methylation machinery (Dnmt1, Dnmt3a and Dnmt3b), Dppa2 and Dppa4 between Dnmt WT (left two columns) and Dnmt TKO (right two columns) following treatment with control siRNA or both Dppa2 and Dppa4 siRNA for 4 days. Hsp90 is shown as loading control. (C) Mass spectrometry quantification of 5-methylcytosine (5mC, left) and 5-hydroxymethylcytosine (5hmC, right) between Dnmt WT (left two columns) and Dnmt TKO (right two columns) cells following control (grey) or Dppa2/4 siRNA (red) treatment for 4 days. Bars represent mean +/-standard deviation of two biological replicates. (D) Expression levels of Dppa2/4-dependent (apricot), -sensitive (green) and -independent (blue) genes between Dnmt WT and Dnmt TKO cells. Data reanalysed from (*25*). (E) Expression of Dppa2 and Dppa4 following control (grey) or Dppa2/4 (red) siRNA treatment in WT (left, dark) and Dnmt TKO (right, light) cells. Relative expression is normalised to the level of control siRNA (dark bars) for WT and DNMT TKO cells. Dots represent biological replicates and bars averages plus standard deviation.

**Supplemental Table 1**

List of all genes, chromosome coordinates, strand, and average log_2_RPM for embryoid body (EB) differentiation experiments. Three independent differentiations were performed (EBset1, EBset2, EBset3) for one wildtype (clone 58) and one Dppa2/4 DKO (clone 43) clone, and samples collected at day 1, 2, 3, 4, 7 and 9 of differentiation.

**Supplemental Table 2**

List of differentially expressed genes (DESeq2+intensity difference filter) for Dppa2-GFP and Dppa4-GFP overexpression lines compared to GFP controls. Columns contain gene name, chromosome coordinates, strand information, classification as differentially expressed in Dppa2 overexpressed cells (Dppa2 O/E, blue), Dppa4-GFP overexpressed cells (Dppa4 O/E, purple), or both Dppa2 and Dppa4 overexpressed cells (Dppa2 O/E and Dppa4 O/E). Log_2_RPM are give for three biological replicates (_1,_2, _3) for two biological clones for GFP (E14+GFP_c1, E14+GFP_c2), Dppa2-GFP (E14+Dppa2_c3, E14+Dppa2_c5) or Dppa4-GFP (E14+Dppa4_c5, E14+Dppa4_c6) overexpressing cells.

**Supplemental Table 3**

qPLEX-RIME results showing Accession, Gene name, description, gene symbol, number of unique peptides detected, peptide intensity for each sample (two biological clones per condition with 3-4 technical replicates each), log_2_ fold change, Average Intensity, t, P-value, adjusted p-value and B value for Dppa2-GFP vs GFP (first tab) versus Dppa4-GFP vs GFP (second tab).

**Supplemental Table 4**

List of Dppa2/4-dependent (first tab), Dppa2/4-sensitive (second tab) and Dppa2/4-independent (third tab) gene promoters (TSS +/-1kb) showing chromosome coordinates, and log_2_ RPM enrichment for Input, Dppa2, Dppa4, H3K4me3, H3K27me3, H2A.Z, Ring1B, Ash2l, Ezh2 and chromosome accessibility for WT and DKO cells. Individual clonal replicates are shown. Input, Dppa2 and Dppa4 data reanalysed from (*9*), all other data from this study.

**Supplemental Table 5**

List of attributes used in machine learning, together with their description and data source (first tab) along with values (second tab).

